# Tropical and montane *Apis cerana* show distinct dance-distance calibration curves

**DOI:** 10.1101/2024.02.10.579752

**Authors:** A. K Bharath Kumar, Ebi Antony George, Axel Brockmann

**Affiliations:** National Centre for Biological Sciences - Tata Institute of Fundamental Research, Bengaluru 560065, India; Department of Apiculture, University of Agricultural Sciences - GKVK, Bengaluru 560065, India; Department of Ecology and Evolution, Biophore, University of Lausanne, 1015 Lausanne, Switzerland

## Abstract

Social bees have evolved sophisticated communication systems to recruit nestmates to newly found food sources. As foraging ranges can vary from a few hundred meters to several kilometers depending on the environment or season, populations living in different climate zones likely show specific adaptations in their recruitment communication. Accordingly, studies in the western honey bee, *Apis mellifera*, demonstrated that temperate populations exhibit shallower dance-calibration curves compared to tropical populations. Here we report the first comparison of calibration curves for three Indian *A. cerana* lineages: the tropical *A. indica*, and the two montane Himalayan populations *A. c. cerana* (Himachal Pradesh) and *A. c. kashmirensis* (Jammu and Kashmir). We found that the colonies of the two montane *A. cerana* populations show dance-distance calibration curves with significantly shallower slopes than the tropical *A. indica*. Next, we transferred *A. c. cerana* colonies to Bangalore (∼ 2600 km away) to obtain calibration curves in the same location as *A. indica*. The common garden experiment confirmed this difference in slopes, implying that the lineages exhibit genetically fixed differences in dance-distance coding. However, the slopes of the calibration curves of the transferred *A. c. cerana* colonies were also significantly higher than those tested in Himachal Pradesh indicating an important effect of the environment. The differences in dance-distance coding between temperate and tropical *A. cerana* lineages resemble those described for *A. mellifera* suggesting that populations of both species independently evolved similar adaptations.

## Introduction

Honey bees use the symbolic dance communication to indicate flight direction and distance of food sources to nestmates in the hive (Frisch, 1967; Riley et al., 2005). This behavior was initially observed in the European honey bee *Apis mellifera carnica* and early on the question was raised whether populations and species of honey bees differ in dance communication similar to dialects in human and bird vocal communication (“dance dialects” see Frisch, 1948). Today there is good evidence that honey bee species and populations of *A. mellifera* differ in several aspects of distance coding and that some of those are heritable (Boch, 1957; Dyer and Seeley, 1991; Johnson et al., 2002; Kohl et al., 2020; Lindauer, 1956; Rinderer and Beaman, 1995; Schneider, 1989). The adaptive tuning hypothesis proposed that the rate of change in dance duration with distance is negatively correlated with the mean foraging range of a population or species (Dyer and Seeley, 1991; Gould, 1982; Kohl et al., 2020; Punchihewa et al., 1985). Thus, populations or species with larger foraging ranges have shallower slopes which reduces the accuracy with which they communicate small changes in flight distances.

The modern honey bee species evolved in the tropics of Asia and only populations of the two cavity nesting species, *A. mellifera* and *A. cerana*, extended their distribution ranges into temperate climate zones (Ji et al., 2020; Ruttner, 1988; Smith, 2020). Compared to the tropics, temperate regions exhibit stronger seasonal variation in food abundance and long periods of food shortage during winter. Colonies of temperate populations are generally larger in worker number and build up substantial food stores (Abrol, 2013; Seeley, 1985). Higher food demands and lower food availability can be countered by enlarging colony foraging ranges (Beekman and Ratnieks, 2000; Couvillon et al., 2014; Grüter and Hayes, 2022; Young et al., 2021). Accordingly, temperate populations of both species show shallower dance-distance calibration curves associated with larger foraging ranges (Boch, 1957; Hu et al., 2023; Kohl et al., 2020; Punchihewa et al., 1985; Sasaki et al., 1993; Schneider, 1989; Su et al., 2008). However, only Boch (1957) compared the dance behavior of different *A. mellifera* populations in common garden experiments to exclude any possible environmental effects. This is crucial as it was later shown that honey bees use optical flow to measure flight distances and use this information for their dance communication (Esch et al., 2001; Srinivasan et al., 2000; Tautz et al., 2004).

In India, there are at least three *A. cerana* lineages according to recent morphological and genomic studies: *Apis indica* (originally called *A. cerana indica* or “Yellow Indian” honey bee), *Apis cerana* (= *A. cerana cerana* or “Black Indian” honey bee), and *Apis cerana kashmirensis* (see Fig 1, Smith, 2020; Abrol, 2013; Hepburn et al., 2001; Qiu et al., 2023; Su et al., 2023). *A. indica* stems from a population that invaded India before the last Ice Age and survived in the southern parts, whereas the *A. cerana* lineage descended from a second colonization (Smith, 2020). Although recent molecular studies support earlier reports that *A. cerana* and *A. indica* might represent separate species, hybrid colonies of black and yellow honey bees are common in the tropical region of southern India (Lo et al., 2010; Oldroyd et al., 2006; Otis and Smith, 2021; Smith, 2020; Su et al., 2023; Viraktamath et al., 2013). Thus, it is unclear whether *A. indica* is a distinct species or an *A. cerana* population at a late state of divergence and we decided to use the term ‘lineage’ to highlight the difference of this bee compared to the other two *A. cerana* populations. The *A. cerana* of the second invasion diverged into several populations adapting to different habitats on the Indian subcontinent. There is good evidence that there are two northern montane populations: *A. c. kashmirensis* in the Jammu & Kashmir region and *A. c. cerana* in the central and eastern Himalayas (Hepburn et al., 2001; Qiu et al., 2023; Su et al., 2023; Viraktamath et al., 2013). These montane bees experience shorter flowering seasons and snowfall for at least a few weeks similar to temperate *A. cerana* and *A. mellifera* populations (Abrol, 2013; Ahmad, 2023; Ji et al., 2020). Both environmental conditions, as discussed above, suggest larger foraging ranges and shallower dance-distance curves compared to the tropical *A. indica*. Starting from this hypothesis we determined the dance-distance calibration curves of *A. c. kashmirensis* (Kashmir), *A. c. cerana* (Himachal Pradesh) and *A. indica* (Bangalore). First, we compared colonies of the two Himalayan lineages, *A. c. kashmirensis* and *A. c. cerana*, within their natural distribution range. Then, we transported three *A. c. cerana* colonies from Himachal Pradesh to the tropical region of Bangalore (∼ 2600 km distance) and performed a common garden experiment testing them in the same location where we had tested colonies of the *A. indica* lineage. We found that colonies of the two Himalayan *A. cerana* populations exhibit dance-distance calibration curves with significantly shallower slopes than those of *A. indica* colonies, and this difference persisted in the common garden experiment. However, the calibration curves of the *A. c. cerana* colonies tested in Bangalore were steeper than those tested in the Himalayas indicating that environmental differences also influence calibration curves. Our findings indicate that temperate and tropical *A. cerana* lineages exhibit similar adaptations in dance-distance coding as temperate and tropical *A. mellifera* populations.

**Fig 1:**
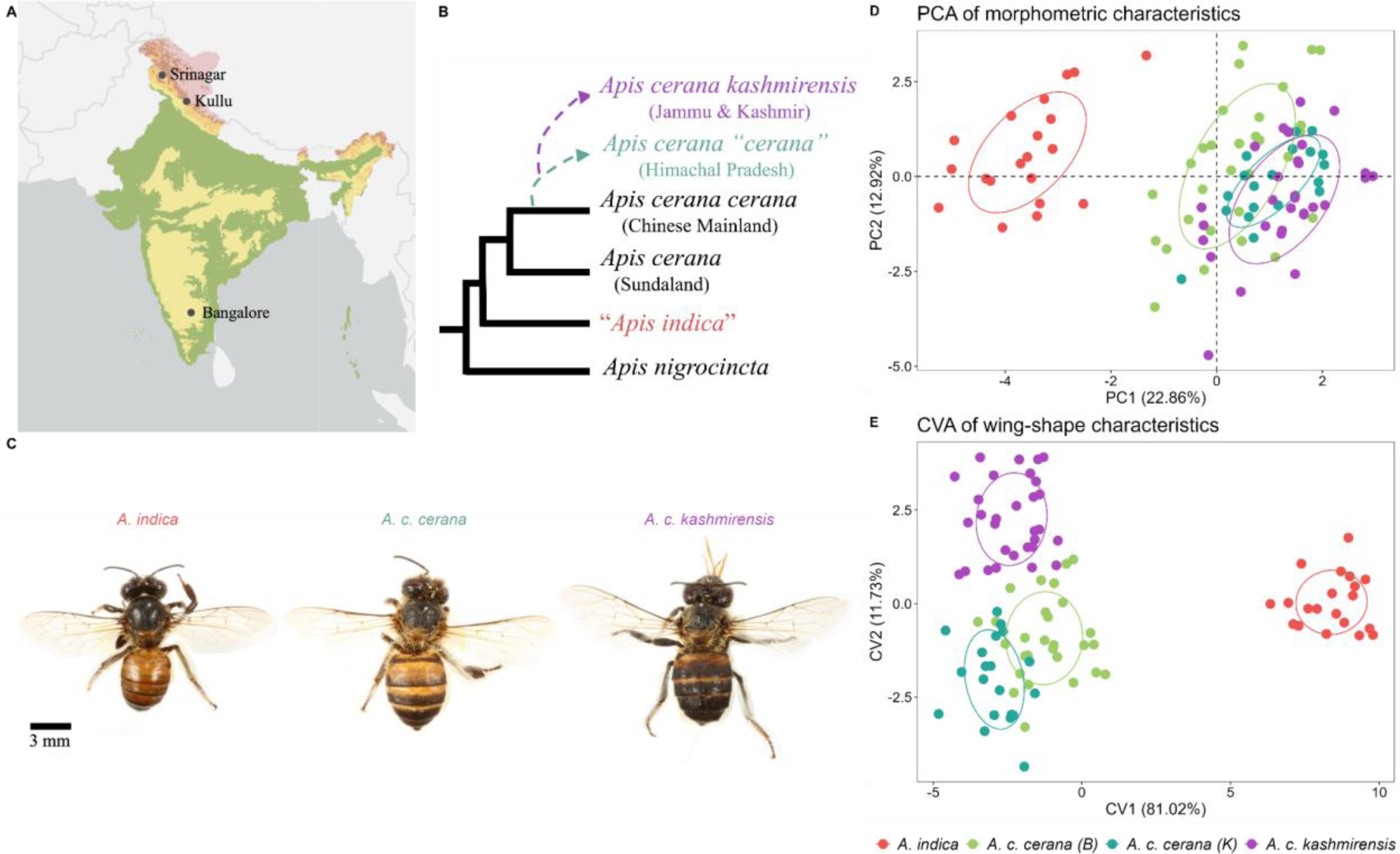
Tropical and montane lineages of *A. cerana* differ in their morphological characteristics. A) Map of India with experimental locations highlighted, B) Phylogeny of the *Apis cerana* group, modified after (Qiu et al., 2023; Smith, 2020) C) Photographs of workers of the three lineages D) PCA biplot of all morphometric characteristics and E) CVA plot of geometric morphometric wing-shape characteristics. In D) and E) the small, filled circles represent individual workers while the large circles represent a normal distribution at the 0.5 level around the points for each lineage in each location. The circles are coloured by lineage and location with red for *A. indica*, green for *A. c. cerana* in Bangalore, teal for *A. c. cerana* in Kullu and purple for *A. c. kashmirensis*.

## Methods

### Honey bee colonies and location of experiments

Colonies of the three *A. cerana* lineages were bought from local beekeepers within their natural ranges: *A. indica* (Department of Apiculture, University of Agricultural Sciences, Bangalore, UAS-GKVK), *A. c. cerana* (Deen Dayal Bee Farm, Kullu, Himachal Pradesh), *A. c. kashmirensis* (Imtiyaz Qureshi Bee Farm, Srinagar, Jammu and Kashmir). In the Himalayas, the *A. c. cerana* and *A. c. kashmirensis* colonies were tested at field sites around 20 km away from the farms of the beekeepers. Three colonies of *A. c. cerana* (obtained from Deen Dayal Bee Farm) were transported by car over a distance of around 2600 km from Kullu to Bangalore at the end of October 2022. They were initially kept on the campus of the National Centre for Biological Sciences. Then the experiments were started at the end of December 2022 on the campus of the neighboring University of Agricultural Sciences, UAS-GKVK. This time schedule was chosen to allow the transported colonies to settle and adapt to the new location and reduce the likelihood of absconding.

### Morphometric measurements

We performed morphometric analyses to confirm the lineage status of the colonies used in this study. From each colony 10 foragers were collected at the end of the experiments described below for standard and geometric morphometric analysis. Photographs of the thorax and wings were taken with a stereomicroscope (Leica M125 C; Camera: Leica MC 190 HD; Software: LAS V4.12). For the photographs the forewings and hindwings were mounted on a glass slide using a coverslip. Measurements were performed on those photographs using Fiji software (Schindelin et al., 2012). For the standard morphometric analysis we measured: the intertegular distance (ID, (Cane, 1987), hamuli number, cubital index (described in Qiu et al., 2023), and fourteen wing characters: forewing length (lfw), forewing width (wfw), length of cubital vein 1 & 2 (Cub1, Cub2), wing vein angles- A4, B4, D7, E9, G18, J16, K19, L13, N23, 026 (Ruttner, 1988).For the geometric morphometric analysis, 20 forewing landmarks were used (Ji et al., 2020). Images of the forewings were first digitized using tpsDig v2.0 and tpsUtil v1.40 (https://sbmorphometrics.org/) and then we obtained the two-dimensional landmark coordinates for further analysis.

### Experimental procedure

We followed the same experimental protocol for all colonies. Table 1 comprises information on colonies, dates, location of experiments and the general flowering conditions during the time the experiments were performed.

**Table 1:**
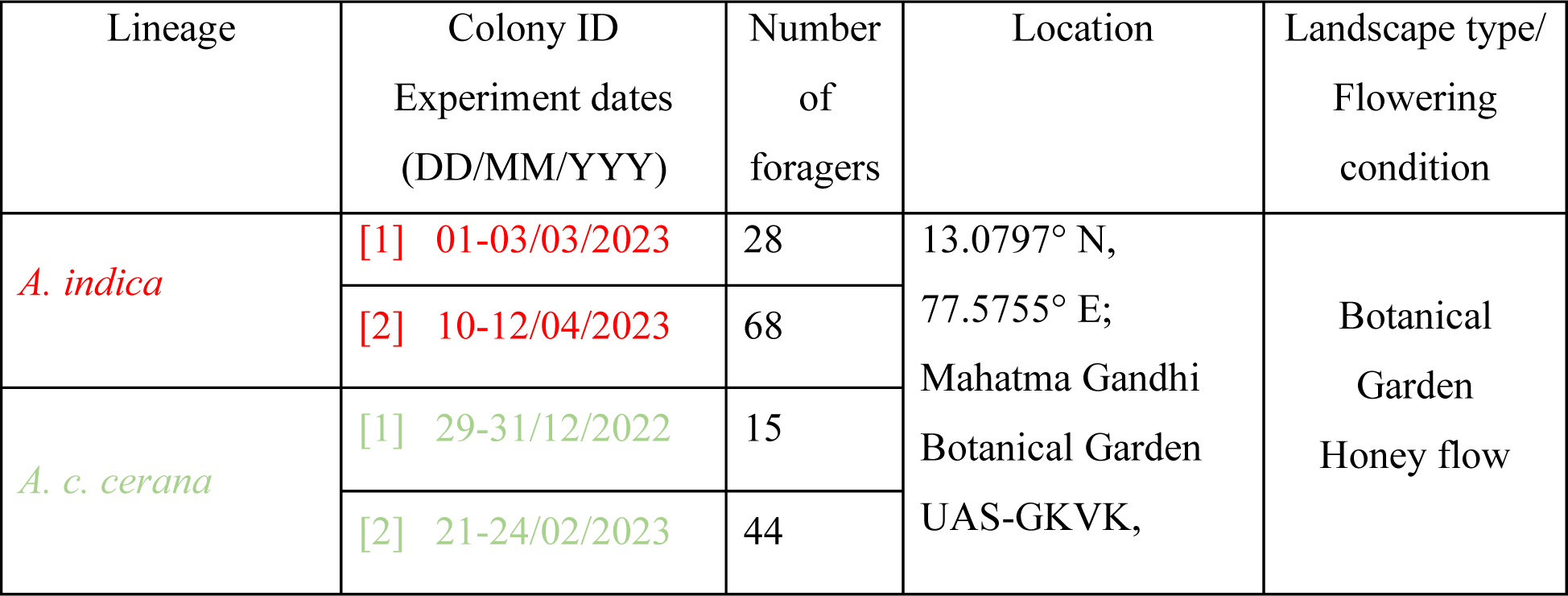
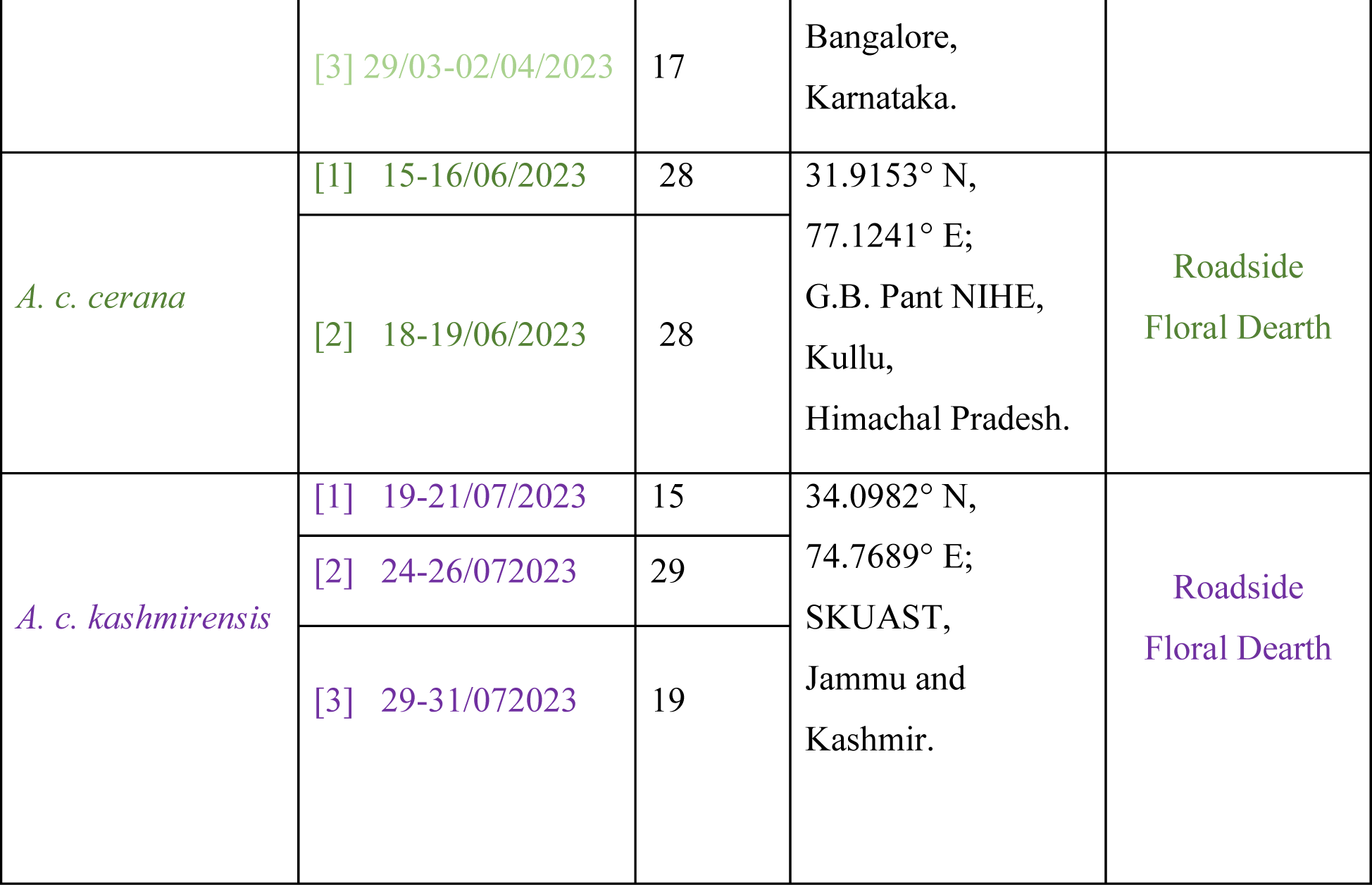
Table of experimental details. The days in which experiments were performed on each colony along with the number of foragers as well the experimental location and the environmental conditions are provided.

Prior to the experiments, colonies were transferred to observation hives during the evening hours after sunset. The observation hives were custom built similar to single hive boxes in which the frames are arranged horizontally, and only one comb was exposed to the observation window (see Fig S1, (Kohl et al., 2020). Colonies were left undisturbed for a day before we started training foragers. To train foragers, a piece of honeycomb filled with 2M sugar syrup was first placed at the entrance of the hive. When a group of about twenty foragers continuously collected sugar syrup, the comb was moved to a gravity feeder containing 2M sugar syrup kept on a stand 10 m from the hive entrance. The piece of comb was removed when all the bees started collecting from the gravity feeder. Bees were individually color-marked on the thorax and abdomen (Uni POSCA Paint Markers, Uni Mitsubishi Pencil, Japan). During the experiments, the feeder setup was gradually moved from one feeder distance to the next at a rate of approximately 10 m per 5 minutes. The first feeder station at which dances were recorded was 50 m from the hive. Feeder stations were separated by 50 m and the bees were trained to a maximum distance of 500 m from the hive in Bangalore and Kashmir, and to a distance of 400m in Kullu (with the exception of colony 1 in *A. c. cerana* Bangalore which could be trained only to 300 m). Waggle dances were recorded using the Sony FDR - AX53 handycam (Sony Corporation, Japan) at 60 frames per second. At each feeder station dances were recorded for 1 hour. Unmarked foragers which arrived at the feeder were collected, kept in boxes during the experiment and released at the end of the day. Experimental duration per colony ranged from 3 to 5 days. Each following day the experiment was resumed at the last feeder position of the previous day.

### Measurement of waggle phases

Analysis of the dance recordings was done using the VitualDub2 software (http://virtualdub2.com/). The waggle phase duration was calculated by counting the number of frames between the start of the waggle phase and the end of the waggle phase and multiplying it with the frame rate of the recording. We defined the start frame as the first frame in which the bee starts to move its abdomen sideways and the end frame as the frame in which the bee stops moving its abdomen back and forth. All the analysis on the waggle dance was performed on the mean waggle phase duration per dance and not on individual waggle phases. Thus, for each distance we had multiple dances from multiple individuals in a given colony.

### Statistics

We used generalized linear mixed models (GLMMs) for our statistical comparisons on the waggle dance. For the morphometric analysis we used dimensionality reduction analysis to identify differences between the four groups as well as linear mixed models for comparing specific traits.

#### Morphometric analysis

Principal Component Analysis (PCA) based on all 17 measured standard morphological traits was used to identify how individual foragers clustered. For the geometric morphometric analysis, Canonical Variate Analysis (CVA) was performed using the size corrected shape data in MorphoJ Ver 1.08.01 (Klingenberg, 2011) to compare differences amongst the lineages. We compared the Mahalanobis distances between the groups and obtained *p*-values from permutation tests (10000 iterations) using MorphoJ.

#### Lineage differences in dance-distance calibration curves

To compare the dance dialects between the different lineages, we fitted linear mixed effects models (LMMs) with the mean waggle phase duration as the response variable, and an interaction between distance (a continuous variable) and lineage (a categorical variable with four levels - *A. indica*, *A. c. cerana* Bangalore, *A. c. cerana* Kullu, and *A. c. kashmirensis*) as the predictor variable. For the random effect structure, we first fit a random slopes model with an effect of colonies on the slope linking mean waggle phase duration and distance. However due to convergence issues with this model we then fit a random intercept model with an effect of colonies on the intercept. After model validation, we used the random intercept model for inference and estimated marginal trends for the slopes for each level of the lineage. We compared significant differences between each pair of slopes using the Tukey method for p-value adjustment.

For comparison of the dance dialects, we also fit non-linear mixed effects models. We did this since previous work had shown that non-linear curves often fit the waggle dance duration-distance relationship better (George et al., 2021; Kohl and Rutschmann, 2021). We fit a logarithmic curve as described previously (George et al., 2021). The relationship between the ‘slopes’ of the non-linear curves fit to each lineage category was qualitatively similar to the results from the LMMs and we report these results in the supplementary material.

#### Colony variation

To look at colony variation in slopes within lineages, we used LMMs. We first created a dummy variable “lineage_colony” which was an interaction between the lineage categorical variable of 4 levels described above and the colony IDs (categorical variable of maximum 3 levels). We then fitted the model with the mean waggle phase duration as the response variable, and an interaction between distance (a continuous variable) and lineage_colony (a categorical variable with ten levels - 2 colonies of *A. indica*, 3 colonies of *A. c. cerana* Bangalore, 2 colonies of *A. c. cerana* Kullu, and 3 colonies of *A. c. kashmirensis*) as the predictor variable. We used this categorical variable to minimize the number of models we ran (4 different models if we looked at each lineage separately instead of 1 model). For the random effect structure we first fit a random slopes model with an effect of individual bees on the slope linking mean waggle phase duration and distance. However due to convergence issues with this model we fit a random intercept model with an effect of individual bees on the intercept. After model validation, we used this model for inference and estimated marginal trends for the slopes for each level of the lineage. We compared significant differences between each pair of slopes using the Tukey method for p-value adjustment. We report differences between colonies of the same lineage in the results.

#### Inter-Individual variation

To look at individual variation in the slope of the relationship between waggle phase duration and distance, we first shortlisted individuals which performed at least one dance at 3 unique distances. This gave us a final data set of 81 individuals across 10 colonies (mean ± sd = 8.1 ± 5.17, range = 1 to 15 individuals). We then create a dummy categorical variable “lineage_colony_beeID” as we described above for the colony comparison incorporating lineage, colony and beeID (our beeIDs were based on colour combinations and hence repeated across colonies and lineages). We then fitted a linear mixed effects model with mean waggle phase duration as the response variable, and an interaction between distance (a continuous variable) and lineage_colony_beeID (a categorical variable with 81 levels for each individual) as the predictor variable. For the random effect structure we first fit a random slopes model with an effect of lineage_colony (described above) on the slope linking mean waggle phase duration and distance. However, due to convergence issues with this model and a random intercept model with an effect of lineage_colony on the intercept, we finally fit a simple linear model without any random effects. After model validation, we used this model for inference and estimated slope values for 81 individuals. We further shortlisted this set of slope values to remove 4 individuals which had negative estimated slopes (either due to low dances in one distance, or very few data points overall).

Next, to compare individual variation across lineages, we obtained the coefficient of variation of individual slopes for each colony within each lineage. Since we had only one individual for *A. c. cerana* Bangalore colony 3, we could not estimate the coefficient of variation for this colony. We then fit a linear model with the coefficient of variation in individual slopes as the response and the lineage (categorical variable of 4 levels) as the predictor. After model validation, we used this model for inference and estimated marginal means for each level of the lineage. We compared significant differences between each pair of estimated marginal means using the Tukey method for p-value adjustment.

We used the R software v4.3.2 (R Core Team, 2023) along with the Rstudio IDE (Posit team, 2023) to perform all the statistical analyses except for the CVA. GLMMs were built using the lme4 package in R (Bates et al., 2015), while we used the nlme (Pinheiro et al., 2023) and aomisc package (Onofri, 2020) to build the non-linear mixed effects models. Model validation was performed using DHARMa (Hartig, 2022) and performance (Lüdecke et al., 2021) packages, while marginal means and contrasts were obtained using the emmeans (Lenth, 2024) and modelbased (Makowski et al., 2020) packages. The PCA was performed using the factoextra package (Kassambara and Mundt, 2020). For the CVA we used the MorphoJ Ver 1.08.01 (Klingenberg, 2011) software. All the plots were created in R using ggplot2 (Wickham, 2016) and cowplot (Wilke, 2024) packages.

## Results

### Morphometric Analysis

The morphological analyses confirmed the phylogenetic lineage of the colonies used in our experiments. The PCA including all the standard morphological characters (i.e. size-dependent and -independent characters) clearly separated the *A. indica* specimens from those of two montane northern lineages *A. c. cerana* and *A. c. kashmirensis* (Fig: 1D). PC1 which accounted for 22.9% of the total variation was mainly influenced by the ID, wfw, lfw, cub1, K19 and ID (loading values of 0.43, 0.42, 0.41, 0.35 and 0.35 respectively), while PC2 which accounted for 12.9% of variation was influenced by cub2, CI, A4 and D7 (loading values of −0.43, −0.41, −0.41 and −0.38 respectively). PC3 which accounted for 11.6% of the variation was influenced by N23, J10, B4, J16 (loading values of 0.46, 0.41, −0.38 and −0.38 respectively. The eigenvalues of PC1, PC2 and PC3 are 4.11, 2.32 and 2.08 respectively and the loading values of all the other variables on each PC are given in Table S1.

A canonical variate analysis (CVA) of geometric morphometric landmarks of the forewings successfully separated the specimens of all three lineages tested (Fig 1E). The Mahalanobis distance between groups were significantly different (*A. indica vs A. c. cerana* Bangalore = 9.94, *p* <0.001; *A. indica* vs *A. c. cerana* Kullu=11.61,p<0.001; *A. indica* vs *A. c. kashmirensis* = 10.98, *p* <0.001; *A. c. cerana* Bangalore vs *A. c. cerana* Kullu =4.05, *p* <0.001; *A. c. cerana* Bangalore vs *A. c. kashmirenis* =3.96, *p* <0.001; *A. c. cerana* Kullu vs *A. c. kashmirenis* =4.55, *p* <0.001). CV1, CV2 and CV3 accounted for 81.02%, 11.73% and 7.23% of the variance and had eigenvalues of 18.74, 2.71 and 1.67 respectively.

### Lineage differences in dance-distance calibration curves

The slopes of mean dance-distance calibration curves based on data from 15-68 foragers per colony and two to three colonies per lineage were significantly different between tropical and montane *A. cerana* lineages (intercept and slope values provided in Fig. 2). The calibration curves of the tropical *A. indica* showed the steepest slope, significantly higher than those of the calibration curves of *A. c. cerana* and *A. c. kashmirensis* (*A. indica* vs *A. c. cerana*, Kullu: difference estimate = 0.0024, t-ratio = 15.09, p < 0.0001; *A. indica* vs *A. c. kashmirensis*, Kashmir: difference estimate = 0.0021, t-ratio = 19.55, p < 0.0001). Moreover, the difference in the slopes between *A. indica* and *A. c. cerana* did not change in the common garden experiment performed in Bangalore (*A. indica* vs *A. c. cerana*, Bangalore: difference estimate = 0.0018, t-ratio = 16.08, p < 0.0001). The slopes of the calibration curves of the two Himalayan lineages *A. c. cerana* and *A. c. kashmirensis* did not differ when tested in the montane regions (*A. c. cerana* Kullu vs *A. c. kashmirensis*: difference estimate = −0.0003, t-ratio = −1.52, p = 0.43). However, *A. c. cerana* colonies transferred from Kullu to Bangalore exhibit calibration curves with significantly higher slopes than the colonies from the same population tested in Kullu (*A. c. cerana*, Bangalore vs *A. c. cerana*, Kullu: difference estimate = 0.0006, t-ratio = 3.73, p = 0.001).

**Fig 2:**
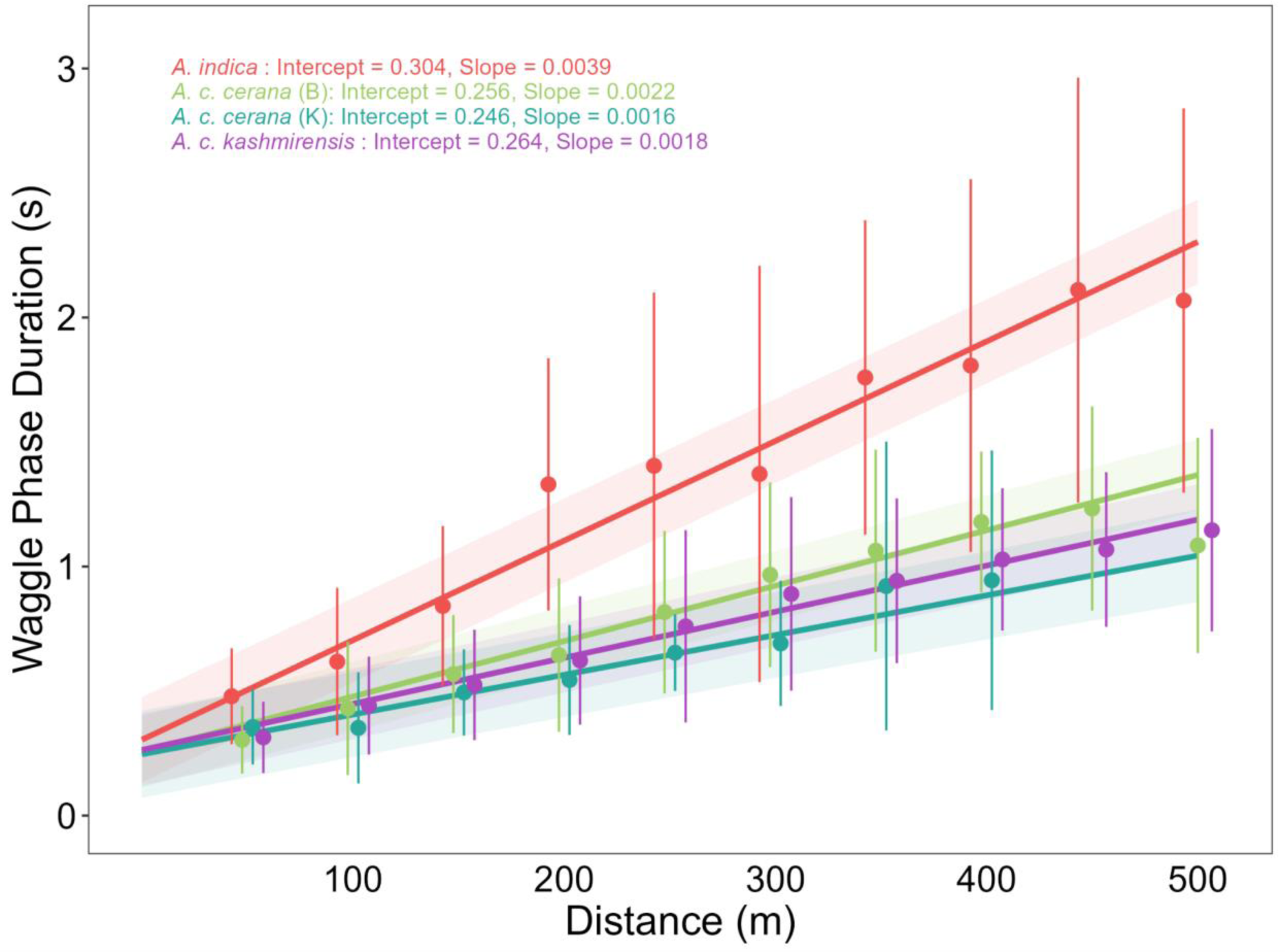
The dance-distance calibration curve of the tropical *A. indica* exhibits a steeper slope than that of the two montane *A. cerana* lineages. Circles and error bars represent mean and standard deviation of waggle phase duration for each distance from 15-68 foragers per colony and 2-3 colonies per lineage in each location. The line represents the estimated trend line obtained from the linear mixed effects model and the shaded region represents the 95% confidence interval around the estimated trend. The inset text provides the intercept and slope values of the estimated trend. Circles, error bars, lines and the shaded region is coloured per lineage and location with red for *A. indica*, green for *A. c. cerana* in Bangalore, teal for *A. c. cerana* in Kullu and purple for *A. c. kashmirensis*.

### Colony variation

Colony variation in the slopes of the calibration curves differed by lineage (slope values provided in Fig. 3). The two *A. indica* colonies significantly differed in the slopes of the dance-distance calibration curves (colony 1 vs colony 2: difference estimate = 0.0011, t-ratio = 7.17, p < 0.0001), whereas the two colonies of *A. c. cerana* as well as the three colonies of *A. c. kashmirensis* tested in their home range did not show significant differences (*A. c. cerana* in Kullu: colony 1 vs colony 2: difference estimate = −0.0006, t-ratio = −1.76, p = 0.762; *A. c. kashmirensis*: colony 1 vs 2: difference estimate = −0.0003, t-ratio = −1.13, p = 0.982; colony 1 vs 3: difference estimate = 0.0001, t-ratio = 0.43, p = 0.999; colony 2 vs 3: difference estimate = 0.0004, t-ratio = 1.50, p = 0.892). In contrast to the experiments in the Himalayas, the three *A. c. cerana* colonies transported and tested in Bangalore showed some degree of variation in the slopes of the calibration curves. Colonies 1 and 2 significantly differed in the slopes, but the slope for colony 3 did not differ from that of colony 1 or 2 (colony 1 vs 2: difference estimate = 0.0014, t-ratio = 3.92, p = 0.004; colony 1 vs 3: difference estimate = 0.0007, t-ratio = 1.57, p = 0.862; colony 2 vs 3: difference estimate = −0.0007, t-ratio = −2.44, p = 0.306).

**Fig. 3:**
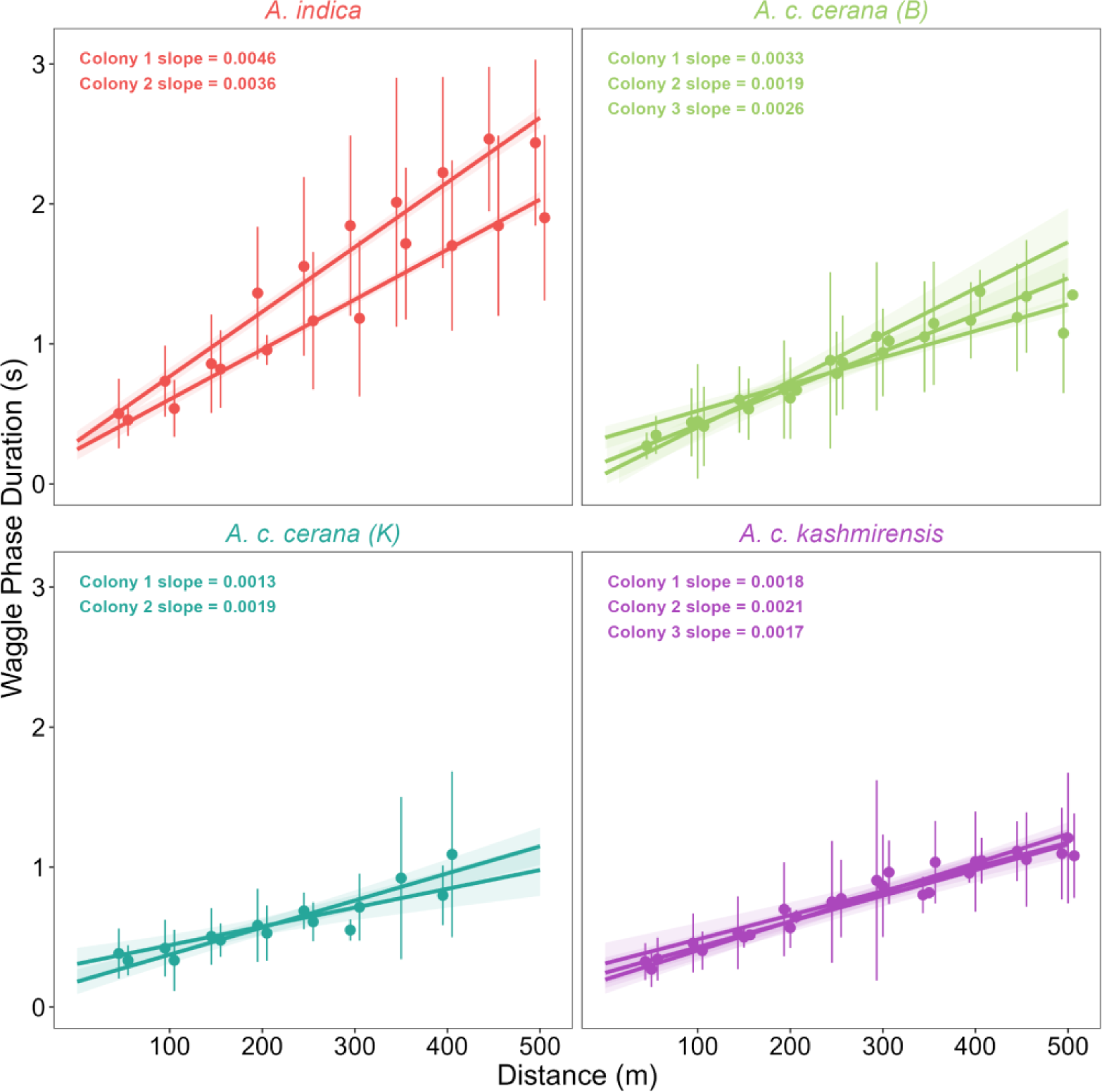
Dance-distance calibration curve of the colonies varies by lineage. Each subplot represents all the colonies tested for one lineage in one location. The circles and error bars represent mean and standard deviation of waggle phase duration for each distance. The line represents the estimated trend line obtained from the linear mixed effects model and the shaded region represents the 95% confidence interval around the estimated trend. The inset text provides the slope values of the estimated trend for each colony. Circles, error bars, lines and the shaded region is coloured per lineage and location with red for *A. indica*, green for *A. c. cerana* in Bangalore, teal for *A. c. cerana* in Kullu and purple for *A. c. kashmirensis*.

### Inter-individual variation

Individual foragers of the different lineages showed considerable variation in the slopes of the dance-distance calibration curves (Fig. 4, and individual slope values provided in Table S2): *A. indica* (minimum slope = 0.0009, maximum slope = 0.0106), *A. c. cerana* in Bangalore (minimum slope = 0.0004, maximum slope = 0.0058), *A. c. cerana* in Kullu (minimum slope = 0.0007, maximum slope = 0.0033), and *A. c. kashmirensis* (minimum slope = 0.0011, maximum slope = 0.0041). Although the variation among individuals was high, the coefficient of variation in the slopes of individuals did not differ amongst lineages and was not affected by environment in *A. c. cerana* (Fig. 4, *A. indica* vs *A. c. cerana*, Bangalore: difference estimate = 0.0908, t-ratio = 0.88, p = 0.817; *A. indica* vs *A. c. cerana*, Kullu: difference estimate = −0.0199, t-ratio = 0.19, p = 0.997; *A. indica* vs *A. c. kashmirensis*: difference estimate = 0.1189, t-ratio = 1.25, p = 0.623; *A. c. cerana*, Bangalore vs *A. c. cerana*, Kullu: difference estimate = −0.1107, t-ratio = −1.07, p = 0.722; *A. c. cerana*, Bangalore vs *A. c. kashmirensis*: difference estimate = 0.0281, t-ratio = 0.29, p = 0.989; *A. c. cerana*, Kullu vs *A. c. kashmirensis*: difference estimate = 0.1388, t-ratio = 1.47, p = 0.516).

**Fig. 4:**
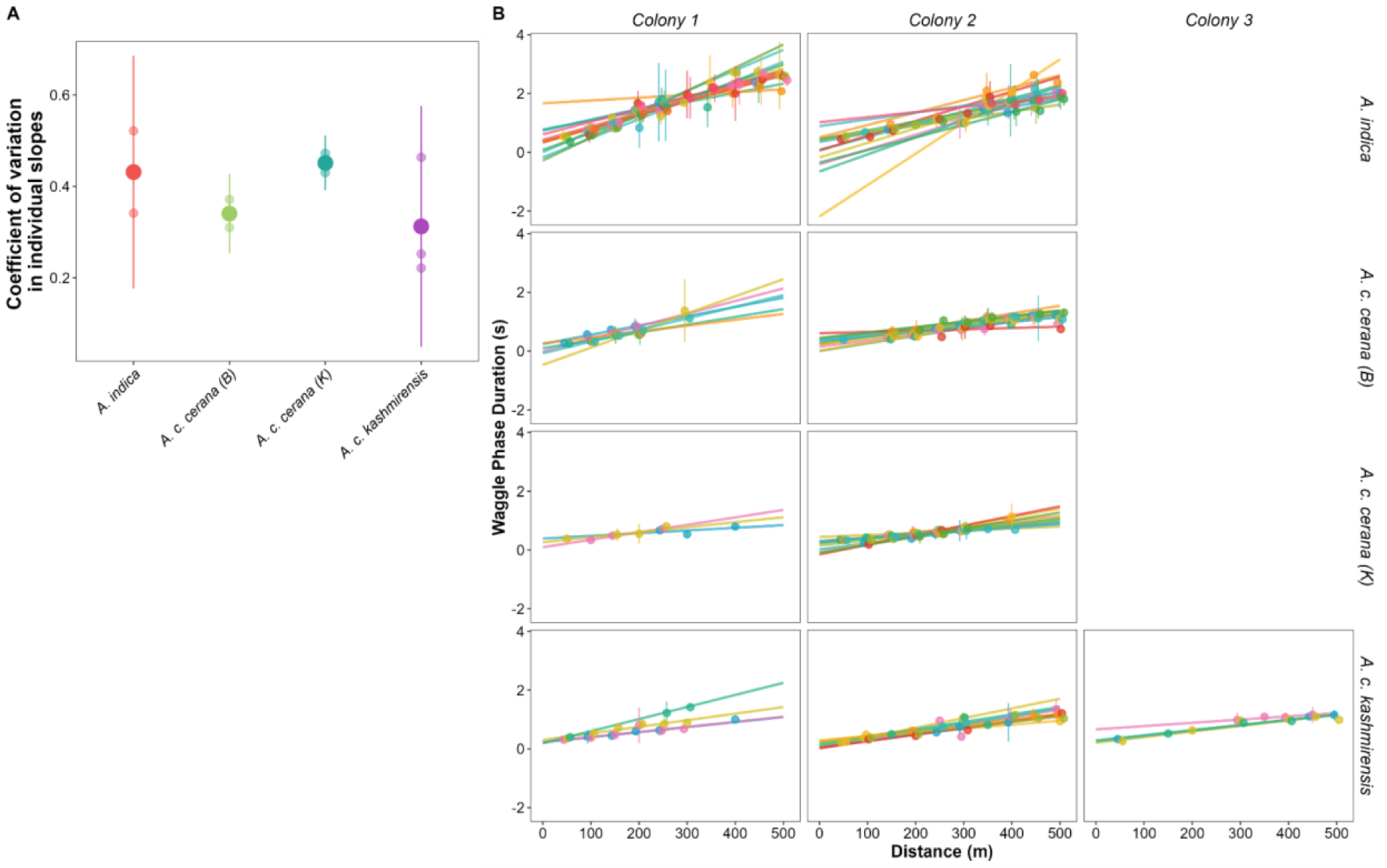
Variation in dance-distance calibration curve of individuals are similar across lineages. A) Small circles represent coefficient of variation of individual slopes for one colony with larger circles and the error bar representing the mean and standard deviation in the coefficient of variation across lineages in each location. Circles and error bars are coloured per lineage and location with red for *A. indica*, green for *A. c. cerana* in Bangalore, teal for *A. c. cerana* in Kullu and purple for *A. c. kashmirensis*. B) Each subplot represents the dance-distance calibration curve for all shortlisted individuals in one colony. Circles and error bars represent the mean waggle phase duration and standard deviation of the same per individual for each distance and the line represents the estimated trend line obtained from the linear model. Circles, error bars and lines are coloured based on individual ID.

## Discussion

Geographic variation in dance behavior (“dance dialects”) was proposed very early in studies on honey bee communication (Boch, 1957; Frisch, 1948), but this idea hasn’t been thoroughly pursued yet, especially in species other than *A. mellifera*. Our study is the first to compare the slopes of dance calibration curves for multiple colonies of different lineages within *A. cerana*. We found that the two montane lineages *A. c. cerana* and *A. c. kasmirensis* exhibit significantly shallower slopes of the curves than the tropical *A. indica*. Furthermore, the difference in the slopes was still present between colonies of *A. c. cerana* examined at the same experimental site where we tested the *A. indica* colonies. However, the slopes of these *A. c. cerana* colonies brought to southern India were also significantly steeper than those of the related colonies tested in the Himalayas. These two results are consistent with the two non-exclusive hypotheses - (i) honey bee populations adapt dance-distance encoding to foraging ranges which depend on environmental and social conditions (Dyer and Seeley, 1991; Kohl et al., 2020), and (ii) dance-distance encoding is not completely innate (Esch et al., 2001; George et al., 2021; Srinivasan et al., 2000; Tautz et al., 2004).

Differences in the distance encoding in dance behavior amongst honey bee species and populations are thought to be a consequence of a limitation in waggle run duration (Dyer, 1991). Populations or species that enlarge their foraging range exhibit shallower dance distance curves in which a unit of dance duration represents a larger flight distance (Dyer and Seeley, 1991; Kohl et al., 2020). Since colonies living in temperate climates experience shorter flowering seasons and have to overcome food shortage during the winter (Abrol, 2013; Seeley, 1985), they likely evolved larger foraging ranges and hence shallower slopes compared to colonies in the tropics. The finding that the temperate lineages of *A. c.* cerana and *A. c. kashmirensis* show calibration curves with shallower slopes compared to the tropical *A. indica* also conforms with the differences in slopes reported between temperate and tropical *A. mellifera* populations (Boch, 1957; Gould and Towne, 1987; Schneider, 1989).

*A. c. cerana* colonies tested in Bangalore not only had higher slopes but higher inter-colony variation than colonies of the same population tested in Kullu. If we assume that all the colonies were genetically similar as they were bought from the same apiary, the question arises why did the variation in dance-distance calibration among colonies increase? As flying insects use optical flow to measure flight distances, generation of dance-distance calibration curves are highly dependent on the visual environment at the experimental site (Esch et al., 2001; Srinivasan et al., 2000; Tautz et al., 2004). So, the difference in the slopes of *A. cerana* colonies tested in Bangalore and Kullu might be a consequence of different optical flow conditions at the different sites. However, an intriguing hypothesis is that the visual optomotor system exhibits some degree of developmental plasticity to facilitate physiological adaptations to variation in visual environments. This could help colonies adapt the dance communication to seasonal variation in flight distances due to changing food availability and foraging ranges. *A. mellifera* colonies surveyed in the UK demonstrated a 5-fold difference in monthly mean foraging distances ranging from 500m in spring to 2500m in autumn (Couvillon et al., 2014; Couvillon et al., 2015). A recent study on honey bee dance behavior showed that dance-distance encoding involves a period of social learning when the bees start foraging (Dong et al., 2023). This finding indicates that there might be a sensitive period that shapes the dance-odometer system. This hypothesis is supported by earlier anatomical studies that demonstrated experience-dependent neuroanatomical plasticity and changes in gene expression in the forager brains during the phase of their orientation flights (Durst et al., 1994; Lutz and Robinson, 2013; Withers et al., 1993).

Furthermore, there is also behavioral evidence that the sun compass system shows a phase of behavioral maturation (Dyer and Dickinson, 1994; Lindauer, 1959). In our study, the two colonies in the Himalayas were tested in two consecutive weeks of flowering dearth, whereas the three colonies in Bangalore were tested over a period of three months with honey flow indicating a high abundance of food sources (personal observations, B.K.). Detailed studies that test for seasonal changes in calibration curves of individual colonies are needed to substantiate the hypothesis that the odometer system might show developmental plasticity.

Our study also provides some insights regarding individual and colony variation in dance distance calibration curves. The level of variation among colonies was different among the lineages and changed with the environment. Colonies of the tropical lineage *A. indica* exhibited the greatest variation, and the *A. c. cerana* colonies tested in Bangalore showed a higher level of variation than the one tested in the Himalayas. In contrast, the variation in dance distance calibration curves among individuals did not differ between lineages. So, tropical lineages did not have greater individual variation than montane lineages. This likely implies that there is a baseline level of variation amongst individuals of a colony. Schürch et al., (2016) examining a total 75 foragers of 3 *A. mellifera mellifera* colonies reported that individual calibrations can vary by as much as 50% independent of environment. The authors suggested that this large variation could arise from their colonies being hybrids of different populations. On the other hand it is also possible that this variation is a consequence of the multiple mating of honey bee queens which generates a high degree of intracolonial genetic diversity (Mattila et al., 2008; Palmer and Oldroyd, 2000; Schürch et al., 2019). Unfortunately, no one has ever tested variation in dance-distance calibration curves among patrilines. Interestingly, there are reports that full sisters preferentially dance with each other, an adequate mechanism to adjust to the variability in the signal that results from genetic diversity (Duong and Schneider, 2008; Oldroyd et al., 1993).

Finally, combining comparative studies of dance behavior in *A. cerana* and *A. mellifera* with population genomics will be a fruitful approach to identify candidate genes involved in distance communication. Both species evolved in the tropics and later extended their distribution range to temperate climate zones, but this happened on different continents, from Africa to Europe in *A. mellifera* and from South-East Asia to Northern Asia in *A. cerana* (Dogantzis et al. 2021; Ji et al., 2020; Smith, 2020). Thus, larger foraging ranges and shallower slopes of the dance-distance calibration curves in temperate populations of *A. cerana* and *A. mellifera* represent parallel adaptations that might have evolved in response to similar selection pressures and is based on similar genomic mechanisms (Gallant et al., 2014; Ji et al., 2020; Jones et al., 2023; Pfenning et al., 2014). In the future, our extensive knowledge of honey bee behavior and the growing availability of genome information and molecular tools for all *Apis* species should increase the attractiveness of this group for comparative evolutionary studies on behavioral variation and the underlying molecular and neural mechanisms (Alves et al., 2023; Bastin et al., 2018; Jourjine and Hoekstra, 2021; Kohl et al., 2020; Seeley 1985).

## Acknowledgments

We are very grateful to Dr. Yeshwanth, H. M. and Mr. Tarun Karmakar, Collection Facility, National Centre for Biological Sciences, Bangalore for providing imaging equipments and help with morphometric analysis; Dr. Jagadish K.S. and faculty of the Department of Apiculture, UAS, GKVK, Bangalore; Dr. A.N. Sringeswara at the Mahatma Gandhi Botanical Garden, GKVK, Bangalore; Dr. Kishor Kumar at G.B. Pant NIHE, Kullu, and Dr. M. A. Paray, Dr Parveena Bano, Dr. Sajad A Ganie for allowing us to perform the experiments; Sajad H. Parey, School of Biosciences and Biotechnology, Rajouri, Jammu and Kashmir, Mr. Khursheed Hussain, Mr. Suriya. S and Mr. Navarasu at SKUAST, Kashmir for helping BK during his research stay. Vidur Niranjan helped in the initial stages of the project and Srihari Hegde managed the transport of *A. c. cerana* colonies from Kullu, Himachal Pradesh to Bangalore by car.

## Author Contributions

Conceptualization: B.K.A.K., A.B.; Methodology: B.K.A.K., A.B.; Formal analysis: B.K.A.K, E.A.G.; Investigation: B.K.A.K.; Writing – original draft: B.K.A.K, E.A.G., A.B.; Writing – review & editing: B.K.A.K, E.A.G., A.B.; Supervision: A.B.; Funding acquisition: A.B.

## Data Availability

Raw data and R scripts used for the analysis is available through Zenodo: https://zenodo.org/doi/10.5281/zenodo.10648086.

