## Supplementary tables for "Tropical and montane *Apis cerana* show distinct dance-distance calibration curves"

**Table S1**

| Measured morphometric characters | | PC1 | PC2 | PC3 |
|---|---|---|---|---|
| Hamuli number | | -0.09265 | -0.02773 | -0.03614 |
| Forewing length (lfw) | | 0.416517 | -0.17188 | -0.02776 |
| Forewing width (wfw) | | 0.423511 | -0.16875 | 0.004622 |
| Length of cubital vein 1 (cub1) | | 0.35961 | 0.177534 | 0.151034 |
| Length of cubital vein 2(cub2) | | -0.18272 | -0.43663 | 0.170353 |
| Cubital index (ci) | | 0.303334 | 0.418637 | -0.07237 |
| Intertegular distance (id) | | 0.435771 | -0.1575 | -0.11641 |
| Wing vein angles | A4 | -0.01281 | -0.41363 | 0.127886 |
| | B4 | 0.134947 | 0.167534 | -0.38582 |
| | D7 | 0.048068 | -0.38493 | -0.12801 |
| | E9 | -0.08611 | 0.112407 | -0.33532 |
| | G18 | 0.096813 | -0.16668 | 0.17138 |
| | J10 | 0.109119 | 0.096257 | 0.410653 |
| | J16 | -0.06391 | 0.124018 | 0.38884 |
| | K19 | 0.353401 | -0.20499 | 0.051394 |
| | L13 | 0.066794 | 0.183381 | 0.143527 |
| | N23 | 0.07931 | 0.174898 | 0.466069 |
| | O26 | -0.06962 | 0.019179 | 0.208748 |

**Table S1: Loading values of the morphometric characters on the first three principal components**

**Table S2**

| Lineage | Location | Colony | Bee ID | Slope | Lower Confidence Limit | Upper Confidence Limit |
|---|---|---|---|---|---|---|
| *A. indica* | Bangalore | 1 | BA_ | 0.0037 | 0.0020 | 0.0054 |
| *A. indica* | Bangalore | 1 | BAI | 0.0044 | 0.0037 | 0.0052 |
| *A. indica* | Bangalore | 1 | GA | 0.0045 | 0.0037 | 0.0053 |
| *A. indica* | Bangalore | 1 | GA_ | 0.0031 | 0.0016 | 0.0046 |
| *A. indica* | Bangalore | 1 | GG | 0.0010 | -0.0022 | 0.0041 |
| *A. indica* | Bangalore | 1 | GT | 0.0070 | 0.0043 | 0.0096 |
| *A. indica* | Bangalore | 1 | RR | 0.0050 | 0.0045 | 0.0056 |
| *A. indica* | Bangalore | 1 | RT | 0.0047 | 0.0042 | 0.0053 |
| *A. indica* | Bangalore | 1 | RY | 0.0048 | 0.0042 | 0.0055 |
| *A. indica* | Bangalore | 1 | WB | 0.0065 | 0.0038 | 0.0092 |
| *A. indica* | Bangalore | 1 | WG | 0.0079 | 0.0045 | 0.0113 |
| *A. indica* | Bangalore | 1 | WT | 0.0043 | 0.0020 | 0.0066 |
| *A. indica* | Bangalore | 1 | WW | 0.0058 | 0.0043 | 0.0073 |
| *A. indica* | Bangalore | 1 | YA | 0.0045 | 0.0041 | 0.0049 |
| *A. indica* | Bangalore | 1 | YY | 0.0046 | 0.0014 | 0.0077 |
| *A. indica* | Bangalore | 2 | B" | 0.0030 | 0.0018 | 0.0041 |
| *A. indica* | Bangalore | 2 | DBT | 0.0050 | 0.0037 | 0.0064 |
| *A. indica* | Bangalore | 2 | Fg! | 0.0049 | 0.0023 | 0.0076 |
| *A. indica* | Bangalore | 2 | FgA | 0.0058 | 0.0026 | 0.0090 |
| *A. indica* | Bangalore | 2 | GA | 0.0042 | 0.0030 | 0.0055 |
| *A. indica* | Bangalore | 2 | GG | 0.0044 | 0.0033 | 0.0055 |
| *A. indica* | Bangalore | 2 | GoT | 0.0107 | 0.0066 | 0.0148 |

| | | | | | | |
|---|---|---|---|---|---|---|
| *A. indica* | Bangalore | 2 | GT | 0.0050 | 0.0044 | 0.0057 |
| *A. indica* | Bangalore | 2 | PA | 0.0024 | 0.0017 | 0.0031 |
| *A. indica* | Bangalore | 2 | RYT | 0.0023 | 0.0002 | 0.0044 |
| *A. indica* | Bangalore | 2 | VT | 0.0030 | 0.0015 | 0.0046 |
| *A. indica* | Bangalore | 2 | Y. | 0.0017 | -0.0002 | 0.0037 |
| *A. indica* | Bangalore | 2 | YY | 0.0043 | 0.0023 | 0.0063 |
| *A. c. cerana* | Bangalore | 1 | BAI | 0.0032 | 0.0015 | 0.0048 |
| *A. c. cerana* | Bangalore | 1 | GT | 0.0043 | 0.0001 | 0.0085 |
| *A. c. cerana* | Bangalore | 1 | OA | 0.0058 | 0.0037 | 0.0080 |
| *A. c. cerana* | Bangalore | 1 | RG | 0.0027 | 0.0006 | 0.0047 |
| *A. c. cerana* | Bangalore | 1 | RT | 0.0020 | -0.0021 | 0.0061 |
| *A. c. cerana* | Bangalore | 1 | RW | 0.0040 | 0.0024 | 0.0055 |
| *A. c. cerana* | Bangalore | 2 | BB | 0.0018 | 0.0006 | 0.0031 |
| *A. c. cerana* | Bangalore | 2 | BT | 0.0021 | 0.0014 | 0.0028 |
| *A. c. cerana* | Bangalore | 2 | RA | 0.0020 | 0.0015 | 0.0025 |
| *A. c. cerana* | Bangalore | 2 | RT | 0.0026 | 0.0014 | 0.0037 |
| *A. c. cerana* | Bangalore | 2 | RW | 0.0026 | 0.0011 | 0.0041 |
| *A. c. cerana* | Bangalore | 2 | RWA | 0.0019 | -0.0004 | 0.0041 |
| *A. c. cerana* | Bangalore | 2 | WA | 0.0027 | 0.0016 | 0.0037 |
| *A. c. cerana* | Bangalore | 2 | WR | 0.0004 | -0.0013 | 0.0022 |
| *A. c. cerana* | Bangalore | 2 | YA | 0.0022 | 0.0010 | 0.0034 |
| *A. c. cerana* | Bangalore | 2 | YT | 0.0016 | 0.0010 | 0.0023 |
| *A. c. cerana* | Bangalore | 2 | YYG | 0.0019 | 0.0010 | 0.0028 |
| *A. c. cerana* | Bangalore | 3 | GP | 0.0028 | 0.0005 | 0.0050 |
| *A. c. cerana* | Kullu | 1 | OT | 0.0009 | -0.0008 | 0.0026 |
| *A. c. cerana* | Kullu | 1 | P* | 0.0025 | -0.0002 | 0.0052 |

| | | | | | | |
|---|---|---|---|---|---|---|
| *A. c. cerana* | Kullu | 1 | WT | 0.0017 | -0.0005 | 0.0039 |
| *A. c. cerana* | Kullu | 2 | BRT | 0.0021 | 0.0008 | 0.0034 |
| *A. c. cerana* | Kullu | 2 | DBT | 0.0012 | -0.0018 | 0.0043 |
| *A. c. cerana* | Kullu | 2 | GRT | 0.0007 | -0.0021 | 0.0036 |
| *A. c. cerana* | Kullu | 2 | GT | 0.0014 | -0.0036 | 0.0063 |
| *A. c. cerana* | Kullu | 2 | LBT | 0.0019 | 0.0009 | 0.0028 |
| *A. c. cerana* | Kullu | 2 | LGT | 0.0016 | 0.0004 | 0.0028 |
| *A. c. cerana* | Kullu | 2 | LGT_ | 0.0030 | 0.0016 | 0.0045 |
| *A. c. cerana* | Kullu | 2 | RA | 0.0033 | 0.0006 | 0.0060 |
| *A. c. cerana* | Kullu | 2 | RBT | 0.0018 | -0.0002 | 0.0038 |
| *A. c. cerana* | Kullu | 2 | WRT | 0.0012 | -0.0004 | 0.0027 |
| *A. c. cerana* | Kullu | 2 | WT | 0.0028 | 0.0001 | 0.0054 |
| *A. c. kashmirensis* | Kashmir | 1 | G/ | 0.0017 | 0.0008 | 0.0026 |
| *A. c. kashmirensis* | Kashmir | 1 | PT | 0.0017 | 0.0007 | 0.0027 |
| *A. c. kashmirensis* | Kashmir | 1 | WA | 0.0022 | 0.0013 | 0.0032 |
| *A. c. kashmirensis* | Kashmir | 1 | YA | 0.0041 | 0.0023 | 0.0059 |
| *A. c. kashmirensis* | Kashmir | 2 | BA | 0.0022 | -0.0001 | 0.0045 |
| *A. c. kashmirensis* | Kashmir | 2 | BB | 0.0027 | 0.0014 | 0.0039 |
| *A. c. kashmirensis* | Kashmir | 2 | BT | 0.0033 | 0.0023 | 0.0043 |
| *A. c. kashmirensis* | Kashmir | 2 | FGI | 0.0025 | 0.0007 | 0.0044 |
| *A. c. kashmirensis* | Kashmir | 2 | OA | 0.0019 | 0.0005 | 0.0032 |
| *A. c. kashmirensis* | Kashmir | 2 | PP | 0.0025 | 0.0002 | 0.0049 |
| *A. c. kashmirensis* | Kashmir | 2 | RWT | 0.0013 | 0.0004 | 0.0023 |
| *A. c. kashmirensis* | Kashmir | 2 | WRT | 0.0023 | 0.0011 | 0.0034 |
| *A. c. kashmirensis* | Kashmir | 2 | YT | 0.0018 | 0.0009 | 0.0027 |
| *A. c. kashmirensis* | Kashmir | 3 | FG_ | 0.0018 | 0.0009 | 0.0028 |

| | | | | | | |
|---|---|---|---|---|---|---|
| *A. c. kashmirensis* | Kashmir | 3 | GWT | 0.0011 | -0.0005 | 0.0027 |
| *A. c. kashmirensis* | Kashmir | 3 | WT | 0.0019 | 0.0012 | 0.0026 |
| *A. c. kashmirensis* | Kashmir | 3 | WW | 0.0018 | -0.0002 | 0.0037 |

**Table S2: Slope values for individual bees.** The table contains information about the estimated slope values of 77 bee IDs and the 95% lower and upper confidence limit around this estimate. The lineages, locations and the colonies individuals belong to are also provided.
